# Functional characterization of ATP13A2 variants associated with distinct neurodegenerative disorders

**DOI:** 10.1101/2023.08.21.552829

**Authors:** Stephanie Vrijsen, Rania Abou El Asrar, Marine Houdou, Chris Van Den Haute, Veerle Baekelandt, Joseph A. Lyons, Jan Eggermont, Peter Vangheluwe

## Abstract

ATP13A2 is a late endolysosomal transporter that exports the polyamines spermine and spermidine from the organellar lumen to the cytosol. Loss-of-function variants in *ATP13A2* are causative for Kufor-Rakeb syndrome (KRS, a recessive juvenile-onset parkinsonism with dementia) and have also been identified in early-onset PD (EOPD) and hereditary spastic paraplegia (HSP). Furthermore, candidate pathogenic ATP13A2 variants have been identified in neuronal ceroid lipofuscinosis (NCL; M854R), multiple system atrophy (MSA; Y1020C) and amyotrophic lateral sclerosis (ALS; I411M) suggesting that ATP13A2 may be implicated in a broader range of neurodegenerative disorders. Since the functional consequences of the NCL, MSA, and ALS variants have not yet been examined, we here characterized these ATP13A2 variants in terms of subcellular localization, cellular polyamine uptake, and transport activity. We found that the homozygous NCL-associated M854R variant results in an instable protein with low expression levels, leading to complete loss of ATPase and cellular polyamine uptake activity. The heterozygous MSA-linked Y1020C variant is properly localized and presents only partially decreased ATPase activity without affecting cellular polyamine uptake. The ALS-associated I411M variant is also correctly localized and exhibits a minor effect on cellular polyamine uptake, however, without a significant impact on ATPase activity. Taken together, only the homozygous NCL variant of ATP13A2 causes a complete loss-of-function, validating that ATP13A2 dysfunction is implicated in NCL. The ALS and MSA variants only presented a subtle functional defect, questioning whether these heterozygous variants are pathogenic and whether ATP13A2 dysfunction may cause MSA or ALS.

## Introduction

The *ATP13A2* (*PARK9*) gene has been genetically implicated in a spectrum of neurodegenerative disorders. First, disruptive mutations in the *ATP13A2* gene were identified in Kufor-Rakeb syndrome (KRS), a severe parkinsonism with juvenile onset and dementia. Since then, over 35 ATP13A2 variants [1,2] have been reported for KRS, early-onset Parkinson’s disease (EOPD), hereditary spastic paraplegia (HSP), neuronal ceroid lipofuscinosis (NCL), amyotrophic lateral sclerosis (ALS), and atypical PD, including multiple system atrophy (MSA) [3-6]. The reported compound heterozygous and homozygous frame shift mutations and early truncations in KRS and HSP clearly points to loss-of-function, which has been biochemically confirmed for a handful of KRS/HSP missense mutations [2].

ATP13A2 belongs to the superfamily of P-type transport ATPases [2], which couple transport of a specific substrate to the hydrolysis of ATP while forming a phospho-intermediate by auto-phosphorylation. ATP13A2 has recently been characterized as a late endolysosomal polyamine transporter that exports endocytosed polyamines from the lumen to the cytosol [7]. Putrescine, spermidine, and spermine are the three main types of polyamines used by mammalian cells, and ATP13A2 presents the highest affinity for spermidine and spermine, the two longest polyamine species [7]. Polyamines are vital for all eukaryotic cells, and are implicated in a broad range of functions. They exert potent chaperone and anti-oxidant activities, but also regulate key processes such as transcription, translation, and autophagy [8]. ATP13A2 presents a neuroprotective role that relies on its polyamine transport function. Indeed, by transporting polyamines ATP13A2 prevents lysosomal polyamine accumulation and subsequent rupture, thereby averting cathepsin B-induced cell death [7]. Polyamines transported by ATP13A2 also counter (mitochondrial-generated) oxidative stress, a hallmark of neurodegeneration [9]. In addition, ATP13A2’s transport function affects α-synuclein membrane association and multimerization [10]. ATP13A2 further displays transport-independent effects, such as maintaining proteostasis by promoting the removal of ubiquitinated proteins via the proteasome [11], and stimulating α-synuclein externalization via extracellular vesicles [10,12], thereby preventing α-synuclein toxicity.

Several ATP13A2 missense variants of uncertain significance have been identified in ALS, MSA, and PD (mainly heterozygous) as well as in HSP, NCL, and KRS (homozygous/compound heterozygous), which are scattered over the coding sequence [1,7,13,14]. Functional data regarding these missense variants of ATP13A2 is incomplete, covering only a limited number of HSP, KRS, and PD mutations with information that is fragmented in literature. Also, the amino acid position of a reported disease variant may refer either to splice variant 1 (1180 amino acid residues, longest isoform and reference sequence) or splice variant 2, which is 5 residues shorter in the N-terminal domain (1175 residues). Splice variant 2 has historically been chosen to investigate the biochemical properties of ATP13A2 and since then has been used as reference protein to assess the impact of mutations. We use the nomenclature according to isoform 1 throughout the manuscript, except for the results section where variants are functionally characterized using isoform 2 (an overview can be found in Table 1). ATP13A2 biochemical activity is measured in overexpression systems by monitoring spermine-dependent ATP hydrolysis or radiolabeled-ATP-dependent auto/de-phosphorylation in membrane fractions as well as by determining cellular uptake of labeled spermine [7,9,15,16]. Typically, a strong correlation between these ATP13A2 activities has been observed.

**Table 1.** Overview of ATP13A2 disease-associated variants used in this study. TM, transmembrane. ^**†**^, mis-labeled in the original reference as M810R.

| Variant isoform 1 | Variant isoform 2 | Disease | Variant status | Variant localization | Reference |
| --- | --- | --- | --- | --- | --- |
| Y1020C<br>c. 3059A>G | Y1015C<br>c. 3044A>G | Multiple System Atrophy | Heterozygous | TM7 | [6] |
| M854R<sup>†</sup><br>c. 2561T>G | M849R<sup>†</sup><br>c. 2546T>G | Neuronal Ceroid Lipofuscinosis | Homozygous | P-domain | [5] |
| I411M<br>c.1233C>G | I406M<br>c. 1218C>G | Amyotrophic Lateral Sclerosis | Homozygous or heterozygous | A-domain | [4] |

In overexpression systems, disease-associated ATP13A2 variants present various combinations of loss-of-function phenotypes, such as mislocalization (T517I, KRS/HSP; Q1140*, HSP; F182L, KRS; G504R, atypical PD), instability (T517I, KRS/HSP; Q1140*, HSP), deficient autophosphorylation (T517I, KRS/HSP; Q1140*, HSP; G877R, KRS) or impaired spermine-dependent dephosphorylation, ATPase or cellular uptake activities (T517I, G877R; KRS/HSP; G533R, T12M, A741T; EOPD) [3,7,17-24]. The emerging view is that KRS-related ATP13A2 variants are more severely impacted as they completely disturb ATP13A2’s transport activity, whereas EOPD-associated variants are relatively mildly affected, which is in line with the severity of the disease symptoms.

So far, the ATP13A2 variants identified in patients with ALS (1 heterozygous and 1 homozygous carrier of the I411M variant [4]), NCL (2 affected homozygous siblings carrying the M854R variant [5]), or MSA (1 heterozygous carrier of the Y1020C variant [6]) have not yet been functionally characterized, which is important for establishing the pathogenicity of the reported variants and extending the spectrum of diseases associated with ATP13A2 dysfunction. We therefore set out to functionally characterize these candidate pathogenic ATP13A2 variants.

## Materials and methods

### Cell culture

Human neuroblastoma SH-SY5Y cell lines either non-transduced (nts) or stably overexpressing WT ATP13A2 or an ATP13A2 variant (D508N, Y1015C, M849R, I406M; using the isoform 2 ATP13A2 protein sequence) were generated via lentiviral transduction as described previously [25]. Cells were maintained at 37 °C in the presence of 5% CO_2_ and incubated in high-glucose Dulbecco’s modified Eagle medium supplemented with 1% penicillin/streptomycin (Sigma), 15% fetal calf serum (heat inactivated) (Sigma), 1% non-essential amino acids (Sigma), 1% sodium pyruvate (Gibco), and selection antibiotic (160 μg/mL hygromycin or 2 μg/mL puromycin (Invivogen)). All treatments were performed in the same medium, but without selection antibiotic.

### Immunofluorescence

Cells were seeded at 75 000 cells/well in a 12-well plate with coverslips. After 48 h, cells were washed with ice-cold PBS, fixed with 4% paraformaldehyde (Affymetrix) for 30 min at 37 °C, and washed again twice with ice-cold PBS. Next, cells were permeabilized and blocked with 5% BSA (Roth) and 0.5% saponin (Sigma) (referred to as blocking buffer) for 1 h. Subsequently, cells were incubated in primary antibody (dissolved in blocking buffer) for 2 h and thereafter subjected to secondary antibody (dissolved in blocking buffer) for 30 min while shaking. Finally, cells were incubated with 200 ng/ml DAPI (D9542, Sigma) for 15 min to visualize the nucleus. In between the different steps, samples were thoroughly washed with PBS. Samples were then mounted and images were acquired using a LSM880 microscope (Zeiss) with a 63x objective and Airyscan detector.

### SDS-page and immunoblotting

Cells were harvested by trypsinization when they reached 70% confluency and subsequently lysed with radio-immunoprecipitation assay buffer (Thermo Fisher Scientific) supplemented with protease inhibitors (Sigma). The protein concentration of the lysate was determined by the use of a bicinchoninic acid (BCA) protein assay. Next, 10 μg of protein was loaded on a NuPage 4-12% Bis-Tris gel (Thermo Fisher Scientific, Bio-Rad) and separated during an electrophoresis run at 150 V in MES running buffer (Thermo Fisher Scientific). Afterwards, the proteins were transferred to a PVDF membrane in NuPage transfer buffer (Thermo Fisher Scientific) supplemented with 10% v/v methanol (Roth). Immunoblots were then incubated in 5% w/v milk powder (1 h, room temperature) for blocking. Next, immunoblots were subjected to primary antibodies that were diluted in 1% w/v BSA O/N (4 °C). After thorough washing, immunoblots were probed with secondary antibody diluted in 1% w/v milk powder (1 h, room temperature). All dilutions and wash steps were performed with TBS supplemented with 0.1% v/v Tween-20 (PanReac AppliChem). Detection was performed by means of chemiluminescence (Bio-Rad ChemiDoc) and protein levels were quantified with Image Lab software.

### BODIPY-spermine uptake

SH-SY5Y cells were seeded at 200 000 cells/well in a 12-well plate. When they reached 70% confluency, the cells were treated for 2 h with 1 μM BODIPY-spermine (37 °C, 5% CO_2_) before collecting them. After centrifugation (450 x g, 5 min), the cells were resuspended in PBS (without Ca^2+^ and Mg^2+^) supplemented with 1% w/v bovine serum albumin (Roth). Next, the mean fluorescence intensity was analyzed of 10 000 events by the use of a FACS Canto II AIG (BD Biosciences) flow cytometer. Data analysis was performed by the use of Flowing Software (created by Perttu Terho, from the Cell Imaging Core of the Turku Centre for Biotechnology).

### ^14^C-labeled polyamine uptake

SH-SY5Y cells were seeded at 200 000 cells/well of a 12-well plate. When cells reached 70% confluency, the experiment was started. Cells were incubated (30 min, 37 °C) either with 5 μM ^14^C-spermine (ARC 3139-50 μCi) or with a mixture of 5 μM ^14^C-spermine and 100 μM unlabeled spermine in medium. Next, medium was aspirated and cells were washed twice with ice-cold PBS (without Ca^2+^ and Mg^2+^). The cells were subsequently lysed by incubation (10 min at room temperature) in 200 μl radio-immunoprecipitation assay buffer (Thermo Fisher Scientific) before scraping. The lysate was added to 7 ml EcoLite Liquid Scintillation Cocktail (MP Biomedicals: 01882475-CF) in scintillation vials. Wells were washed once with 200 μl ice-cold PBS (without Ca^2+^ and Mg^2+^), which was also added to the accompanying scintillation vial. After mixing the vials, ^14^C radioactivity in counts-per-minute (CPM) was measured with liquid scintillation counting (TRI-CARB 4910TR V Liquid Scintillation Counter, PerkinElmer).

### Microsome collection

SH-SY5Y cells overexpressing WT ATP13A2 or an ATP13A2 variant were seeded in 500 cm^2^ plates. Once they reached 70-80% confluency, cells were collected. Next, cells were lysed by resuspending the cell pellet in 3 ml hypotonic LIS buffer (10 mM Tris.HCl pH 7.5, 0.5 mM MgCl_2_.6H_2_O, 1 mM DTT, 1x SigmaFast protease inhibitors), which was incubated on ice for 15 min. The suspension was transferred to a Dounce homogenizer and 60 up-and-down strokes were applied, followed by addition of 3 ml 1 M solution (0.5 M sucrose, 10 mM Tris.HCl pH 7.3, 40 μM CaCl_2_, 1 mM DTT, 1x SigmaFast protease inhibitors) and another 30 up-and-down strokes. The nuclear (1000 x g, 10 min, 4 °C), mitochondrial-lysosomal (12 000 x g, 20 min, 4 °C), and microsomal fractions (140000 x g, 35 min, 4 °C) were then collected. Fractions were suspended in 0.25 M sucrose with 1x SigmaFast protease inhibitors.

### ADP-Glo assay

ATPase activity was assessed using a commercially available luminescence assay (ADP-Glo Max assay, Promega) according to manufacturer’s instructions. A reaction mixture (final volume 25 μl) was made in a 96-well plate and contained 50 mM MOPS-KOH (pH 7), 100 mM KCl, 11 mM MgCl_2_, 1 mM DTT, 0.1 mg/ml DDM, 5 μg microsomes (1:2 ratio DDM: microsomes) and various concentrations of polyamines. The microsomes were collected from SH-SY5Y cells overexpressing ATP13A2 (WT or variants). Next, the reaction was incubated (20 min, 37 °C followed by 10 min, RT) following addition of 5 mM ATP and terminated by adding 25 μl of ADP-Glo Reagent. The 96-well plate was subsequently incubated (40 min, room temperature), followed by the addition of 50 μl of ADP-Glo Detection Reagent. After 60 min, luminescence was detected using the Bio Tek plate reader.

## Results

Via transduction with different viral vector dilutions, we generated SH-SY5Y cell lines overexpressing different levels of the candidate pathogenic ATP13A2 variants I406M, M849R, and Y1015C (Table 1; residue numbers correspond to ATP13A2 splice variant 2) relative to ATP13A2 WT and a catalytically dead variant (D508N). We then assessed the expression levels of WT *versus* mutant proteins to select stable cell lines expressing comparable protein levels. The expression level of the M849R variant was significantly lower, even at the highest viral particle load, indicating that this protein may present a decreased stability (Fig. 1A). Next, we assessed the localization of the variants with immunocytochemistry and found that similar to WT, the MSA and ALS variants colocalized with the late endolysosomal marker CD63, whereas the M849R variant was localized in a mesh-like network (Fig. 2), suggesting that it may be trapped in the endoplasmic reticulum.

**Fig. 1.**
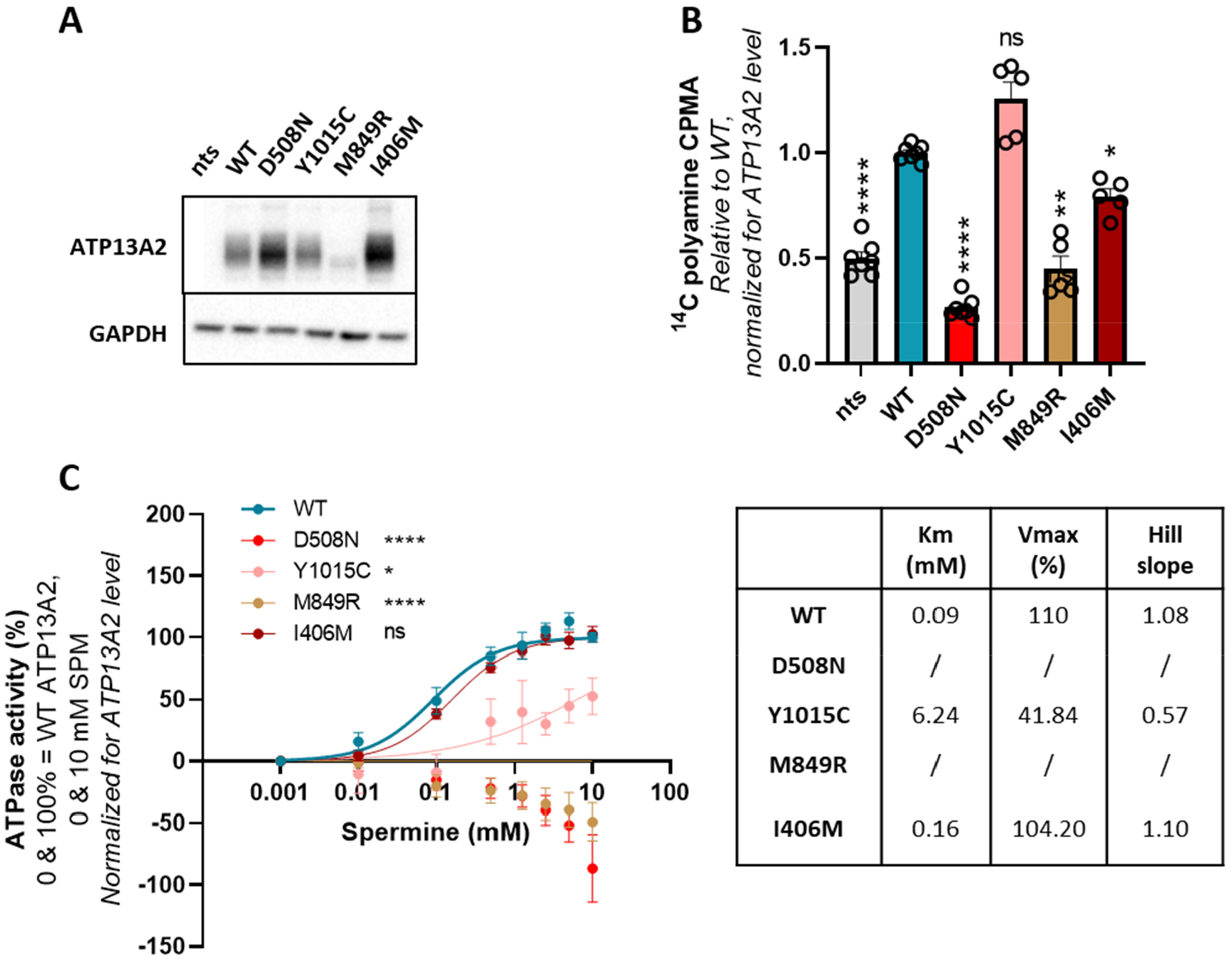
Characterization of ATP13A2 disease-associated variants. **(A)** Lysates were collected and subjected to immunoblotting for SH-SY5Y neuroblastoma cells with endogenous ATP13A2 levels (non-transduced, nts) or stably overexpressing wild-type ATP13A2 (WT), a catalytically dead variant (D508N) or a disease-associated variant (Y1015C, M849R, or I406M). A representative Western blot is shown. **(B)** Polyamine uptake activity was measured with a ^14^C-radiolabeled spermine uptake assay. **(C)** The ATPase activity was determined by the use of isolated microsomes. The table lists the Km, Vmax and Hill slope values for every variant. Data are the mean of a minimum of three independent experiments ± SEM. Data in panel **(B, C)** are normalized for ATP13A2 protein level if the protein is expressed. CPMA, Counts Per Minute. * p<0.05; ** p<0.01; **** p<0.0001; ns, non-significant *versus* WT; ANOVA post-hoc Tukey’s multiple comparison test.

**Fig. 2.**
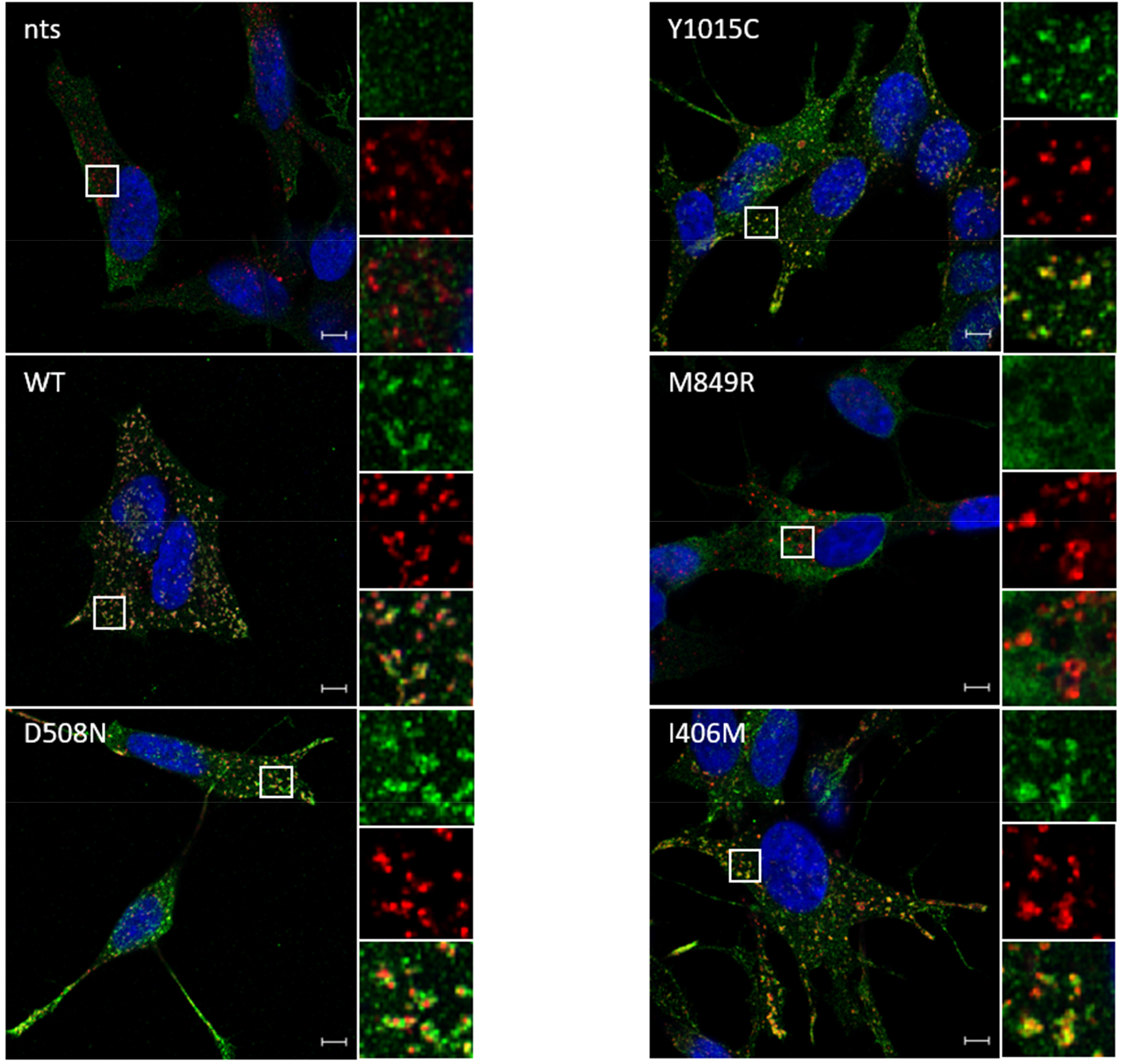
Subcellular localization of ATP13A2 disease-associated variants. SH-SY5Y neuroblastoma cells with endogenous ATP13A2 levels (non-transduced, nts) or stably overexpressing wild-type ATP13A2 (WT), a catalytically dead variant (D508N) or a disease-associated variant (Y1015C, M849R, or I406M) were subjected to immunofluorescence to assess ATP13A2 (in green) colocalization with the late endolysosomal marker CD63 (in red). Scale bar, 5 μm. Images are representative for n=3 experiments.

To assess the functional impact of the mutations, we performed a ^14^C-radiolabeled spermine uptake assay with intact cells as a measure of ATP13A2-mediated polyamine uptake activity and we conducted an ADP-Glo assay on isolated microsomes to measure spermine-dependent ATPase activity. Similar to the catalytically dead D508N variant, the M849R variant exhibited a significantly lower activity in both assays, clearly demonstrating loss-of-function (Fig. 1B, 1C). The functional impact of the ALS and MSA variants is more modest. Whereas cellular ^14^C-radiolabeled spermine uptake by the ALS I406M variant is decreased by approximately 20%, this is not corroborated by the ATPase assay in microsomes (Fig. 1B, 1C). The Y1015C MSA variant presents a decreased ATPase activity in microsomes, but surprisingly, without significantly impacting the ^14^C-spermine uptake in intact cells (Fig. 1B, 1C). We therefore decided to repeat the cellular polyamine uptake assay using previously validated fluorescently-labeled [7,9,16,26] BODIPY-spermine (Fig. S1), which confirmed our results obtained with radiolabeled spermine. Together, our data point to distinct functional phenotypes of the reported NCL, ALS, and MSA-associated missense variants of ATP13A2.

## Discussion

To evaluate whether ATP13A2 may be involved in other neurodegenerative disorders than KRS, HSP, and PD, we here biochemically examined the functional impact of three uncharacterized ATP13A2 variants reported in patients with NCL, ALS, and MSA.

The homozygous ATP13A2 variant identified in two siblings with NCL (M854R, isoform 1; M849, isoform 2; mis-labeled in the original reference as M810R) [5] appears mis-localized and presents low expression indicating that it may be unstable. This at least partially explains the lower cellular spermine uptake in cells as well as the decreased ATPase activity in isolated microsomes *versus* WT ATP13A2, demonstrating that this represents a severe loss-of-function variant. These results provide strong functional support for the genetic association of *ATP13A2* mutations with NCL, a neurodegenerative lysosomal storage disorder marked by the accumulation of lipofuscin. This aligns with observations in animals, since *Atp13a2* knockout mice accumulate lipofuscin in the cerebellum, hippocampus, and cortex [27], whereas Tibetan terriers manifest a late-onset lethal form of NCL as a result of a single base pair deletion (c.1620delG) within exon 16 of the *ATP13A2* gene resulting in an ATP13A2 protein that lacks 69 amino acid residues [28]. An ATP13A2 missense variant was also identified in Australian Cattle Dogs with late onset neuronal ceroid lipofuscinosis [29].

We also studied a heterozygous ATP13A2 variant (Y1020C, isoform 1; Y1015C, isoform 2) identified in a 54-year old Italian patient with MSA [6], a severe neurodegenerative α-synucleinopathy. The patient presented gait and speech difficulties associated with orthostatic dizziness and urinary incontinence, and was initially levodopa responsive. Brain MRI demonstrated suggestive MSA features, including putaminal atrophy and hot cross bun sign with pontocerebellar atrophy [6]. Here, we report that the Y1020C variant presents a reduced ATPase activity without significantly affecting cellular polyamine uptake, questioning the functional impact of this variant in the heterozygous state. Further screening of an MSA cohort of 100 patients identified two other missense variants (A249V and R294Q) on positions that are highly conserved, as well as seven synonymous variants [6]. However, no functional evidence has been provided to demonstrate causality. The heterozygous A249V and R294Q variants have been reported before in two patients with EOPD, although the R294Q variant was also found in a healthy control [30]. Establishing the link between ATP13A2 and MSA will therefore require additional functional, genetic, and clinical confirmation.

ALS patients exhibit a higher prevalence of Lewy body disease and 30% of ALS patients present parkinsonian features. PD-causative genes have therefore been proposed as risk modifiers in ALS and a higher frequency of mutations in autosomal recessive PD genes has been reported in the ALS *versus* control cohort [31]. The genetic link between ATP13A2 and ALS is however not solid, and is mainly based on a report describing a patient carrying a homozygous truncation variant in ATP13A2 (among variants in more than 25 other genes) [4]. The authors specified variant E613* terminating the N-domain of ATP13A2, which would completely disrupt transport activity. The authors further reported that the E613* ATP13A2 truncation variant was unable to restore cerebellum and motor neuron defects in a zebrafish model of atp13a2 knockdown [4]. However, the residue number E613 must be incorrect, since no Glu residue can be found at this position of splice variant 1 or 2; and we were unable to verify the sequencing data questioning these results. Moreover, the patient with the truncation mutation was first diagnosed with complex HSP based on symptoms involving the lower limbs. The diagnosis was later revised to a complex phenotype with prominent ALS-like characteristics, such as the presence of symptoms in both upper and lower motor neurons, tongue atrophy with fasciculations and mild intellectual disability [4]. Additional screening for *ATP13A2* mutations in genetic ALS datasets performed within the same study only revealed one other homozygous variant of *ATP13A2* within the ALS cohort (I411M in isoform 1 or I406M in isoform 2) [4], which we here functionally assessed. We found that this variant had only a modest impact on spermine uptake in cells (a decrease of approximately 20%), while the ATPase activity in microsomes was not affected, questioning whether this variant may be disease-causing. Four more variants in *ATP13A2* have recently been identified in a cohort of 330 ALS patients, but detailed clinical and allele information is lacking [31]. Of these, the A461Vfs*5 variant is most likely disruptive, but was identified in a patient with disease onset at the age of 87. No information is available for the P983S variant, whereas the I946F and Y1020C variants have also been observed in a patient with either PD [32] or MSA [6], respectively. The ALS patient with the Y1020C variant also carried a mutation in *SOD1*, a known ALS-causative gene [31]. It is clear that additional genetic, clinical, and biochemical evidence will be required before a genetic association of ATP13A2 with ALS can be conclusively established.

The observed discrepancy between *in vitro* and *in cellulo* data for the I411M and Y1020C variants may depend on factors that are only present in intact cells and may be lacking in the membrane fractions. Although we were unable to identify a clear loss-of-function effect of the MSA and ALS variants on the polyamine transport function of ATP13A2, other factors may mask a loss-of-function phenotype, such as a compensatory upregulation of (an)other polyamine transporter(s) or other regulatory elements. We further cannot rule out that the mutations affect the transport-independent scaffold function of ATP13A2 [11] or the responsiveness of the ATP13A2 transport activity to (unknown) regulatory factors or stress conditions in cells. We have evaluated ATP13A2 activity in basal conditions, but ATP13A2 activity may become limiting when other genetic and/or environmental insults are present, such as α-synuclein toxicity, oxidative stress, and/or mitochondrial-lysosomal dysfunction. Indeed, ATP13A2-deficient SH-SY5Y cells show increased sensitivity for rotenone and other oxidative stress inducers [9], while its expression is under control of the hypoxia-inducible transcription factor HIF1α [33], indicating that this transporter may be crucial in stress conditions.

Finally, several disease variants (EOPD, MSA, ALS) have been reported in the heterozygous state, but so far, their functional impact appears relatively modest (MSA and ALS, this study; EOPD [7]), questioning whether they may exert any dominant negative effects that affect the wildtype allele, which will be examined in future studies.

## Conclusion

In this study, we have conducted a comprehensive biochemical characterization of ATP13A2 variants linked to NCL (M854R), MSA (Y1020C), and ALS (I411M). Only the NCL variant stands out due to its pronounced loss-of-function phenotype, providing compelling support for the established genetic link between ATP13A2 and NCL. Nonetheless, in light of the moderate functional influence of the heterozygous ALS and MSA variants studied so far, there is a need for additional biochemical, genetic, and clinical investigations to conclusively establish the genetic association between ATP13A2 and ALS or MSA.

## Funding and acknowledgements

The authors acknowledge the financial support of KU Leuven Consortium InterAction (C15/15/073 to P.V.) and the Fonds Wetenschappelijk Onderzoek (FWO) Flanders (G094219N to P.V. and 1S88419N to S.V.). P.V., R.A.E.A., and M.H. are funded by the joint efforts of The Michael J. Fox Foundation for Parkinson’s Research (MJFF) and the Aligning Science Across Parkinson’s (ASAP) initiative. MJFF administers the grant ASAP-000458 on behalf of ASAP and itself.

## Disclosure statement

P.V. is recipient of a Michael J. Fox Foundation for Parkinson’s Research grant and works together with an industrial partner to identify and characterize small molecule activators of ATP13A2 for Parkinson’s disease therapy. The authors are not aware of any other affiliations that might be perceived as affecting the objectivity of this manuscript.

**Fig. S1.**
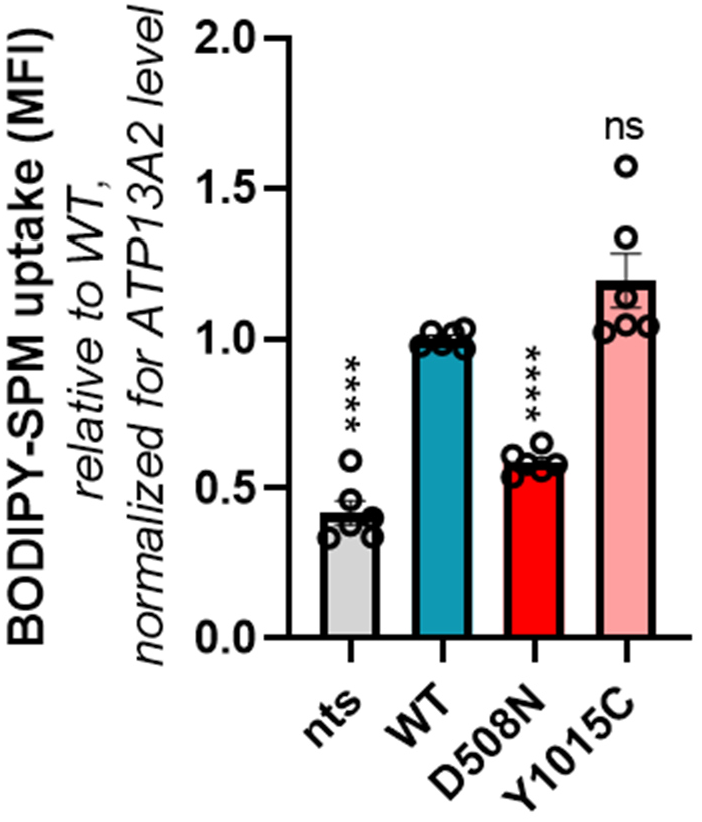
The Y1015C ATP13A2 variant does not alter cellular polyamine uptake. SH-SY5Y neuroblastoma cells with endogenous ATP13A2 levels (non-transduced, nts) or stably overexpressing wild-type ATP13A2 (WT), a catalytically dead variant (D508N), or the Y1015C disease-associated variant were exposed to 1 μM BODIPY-spermine (BODIPY-SPM) for 2 h to assess polyamine uptake. Data are the mean of a minimum of three independent experiments ± SEM. Data are normalized for ATP13A2 protein level if the protein is expressed. MFI, Mean Fluorescence Intensity. **** p<0.0001; ns, non-significant *versus* WT; ANOVA post-hoc Tukey’s multiple comparison test.

